# Effort experience as a driver of behavioral modulation in *Caenorhabditis elegans*

**DOI:** 10.64898/2026.10.09.758103

**Authors:** Reema Arbid, Ithai Rabinowitch

## Abstract

Effort is typically studied as a cost that influences choice, motivation, or persistence, but less is known about whether effort embedded within ordinary behavior can itself leave a trace that alters subsequent behavior. We developed a *Caenorhabditis elegans* paradigm in which animals reached the same food source under low or high effort conditions and were then tested in various behavioral contexts. Prior high effort enhanced subsequent food approach, generalized to more demanding tests including aversive barrier crossing, and persisted for at least 1 h but was lost by 24 h. The effect was distinct from food deprivation and was accompanied by context-dependent changes in locomotor dispersal rather than a uniform increase in activity. A monoamine screen implicated tyramine signaling. High effort also reduced ongoing activity in the tyraminergic RIM interneurons and enhanced the AWC OFF sensory response. These findings identify effort as a behaviorally consequential component of ordinary experience and suggest that ongoing behavior can generate internal state changes that may recursively influence future behavior.

## Introduction

Experience-dependent behavioral plasticity allows prior experience to shape future behavior. This is most evident in dedicated learning paradigms, including classical conditioning, habituation, and reinforcement learning, in which particular stimuli, actions, or outcomes acquire altered significance^1–3^. Beyond such separable episodes of learning, everyday behavior also varies from moment to moment in how it is performed. For example, the same action may require less or more sustained movement, time, or energetic investment depending on circumstances. Whether such variation within ordinary behavior itself leaves a trace that subsequently modifies behavior is less clear.

Effort is a prominent feature of experience that may produce such carryover. Previous work has shown that high effort can influence subsequent persistence and performance^4,5^, and can also alter the valuation of outcomes and choices between alternatives^6,7^. Accordingly, effort has been studied both as an imposed demand and as a cost incorporated into decision making, where it interacts with reinforcement, reward magnitude, motivation, and action selection^8,9^ . Recently, effort-based choice has also been demonstrated in *C. elegans*, which discounts food options according to the effort required to obtain them^10^. This establishes that even the compact nematode nervous system can incorporate effort costs into behavioral decisions. Whereas that work addressed effort prospectively, as a cost influencing choice, we examine effort retrospectively, as an experienced feature of behavior that may modify what the animal does next.

To isolate this question, we developed a paradigm in which *C. elegans* ultimately reached the same food source but differed in the amount of sustained action required to do so. Thus, effort was manipulated as a property of the experience itself, while avoiding a choice between alternatives and minimizing differences in the eventual outcome. This allowed us to ask whether effort embedded within an otherwise routine action can become behaviorally consequential after that action has ended.

*C. elegans* provides a useful system for probing how such experience is translated into subsequent behavioral modulation. Its compact nervous system, defined neuromodulatory pathways, genetic accessibility, and optical access to identified neurons make it possible to connect behavioral modulation to specific molecular and cellular processes. Using behavioral assays, monoamine-pathway mutants, and calcium imaging, we find that effort during food approach leaves a persistent, context-dependent behavioral trace, implicates tyramine signaling, and is accompanied by altered activity in RIM and AWC neurons.

## Results

### Effortful food approach enhances subsequent food approach

Effort is often associated with conspicuously demanding activities, such as strenuous exercise or heavy physical work. But effort also pervades ordinary behavior, to a smaller and more variable degree, from reaching for an object on a high shelf to making an additional stop along a route. Since experimentally imposed high effort is known to alter subsequent behavior^4,5^, we asked whether elevated effort embedded within an otherwise ordinary behavior may similarly leave a trace reflected in later behavioral modulation.

To address this question, we developed a simple *C. elegans* paradigm in which animals experienced either low or high effort while accessing food (Fig. 1a). In the low effort condition, worms were placed directly on a bacterial food patch. In the high effort condition, worms were placed at the opposite side of the plate and had to traverse the agar surface to reach the same food source. After 1.5 h, animals from both groups were transferred to identical test plates, placed opposite a food target, and after 20 min the number of worms on the half plate containing the food target was counted as well as the other half, and a target arrival index was derived (Fig. 1a). Animals previously experiencing high effort subsequently accumulated more strongly at the food target than animals from the low effort condition (Fig. 1b). Thus, even a relatively simple episode of effort during food approach was sufficient to alter subsequent food-approach.

**Figure 1.**
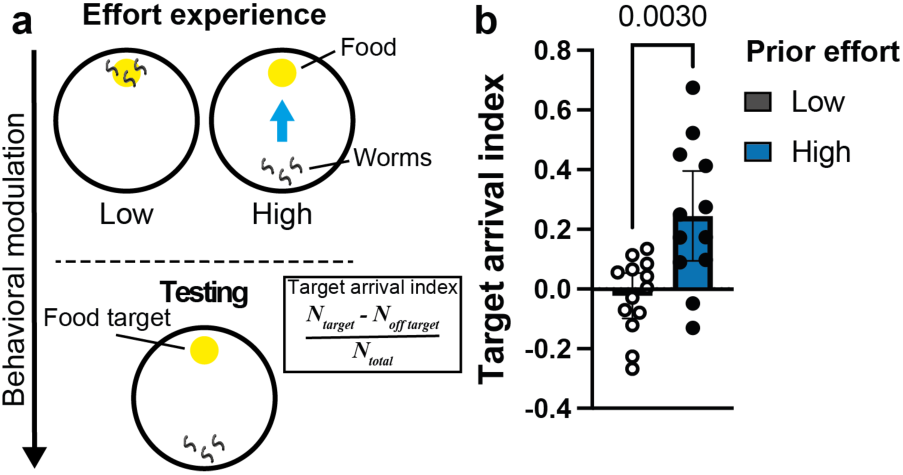
Effortful approach to food enhances subsequent food approach. (**a**) Schematic of the effort experience and subsequent food-approach test. (**b**) Target arrival following low or high effort experience. Welch’s *t*-test; *n* = 13, 12 (low, high). Each point represents an independent experimental replicate (test plate). Bars show mean ± 95% CI.

### Food deprivation does not account for effort-induced behavioral modulation

A salient difference between the low and high effort conditions is the time spent without food. Whereas low effort animals are placed directly on food, high effort animals remain food deprived while traversing the plate and only encounter food upon reaching the target. We therefore asked whether food deprivation itself could account for the subsequent behavioral modulation. To test this possibility, we repeated the experiment, this time including a no-food condition in which animals were placed on plates without a bacterial patch for the same 1.5 h duration (Fig. 2a). If food deprivation were to underlie the effect, then prolonged deprivation would be expected to mimic, or potentially enhance, the behavioral change observed after high effort. Instead, no food animals showed a target arrival index comparable to low effort animals, significantly lower than that of high effort animals (Fig. 2b), indicating that food deprivation alone does not reproduce the effect of effort experience.

**Figure 2.**
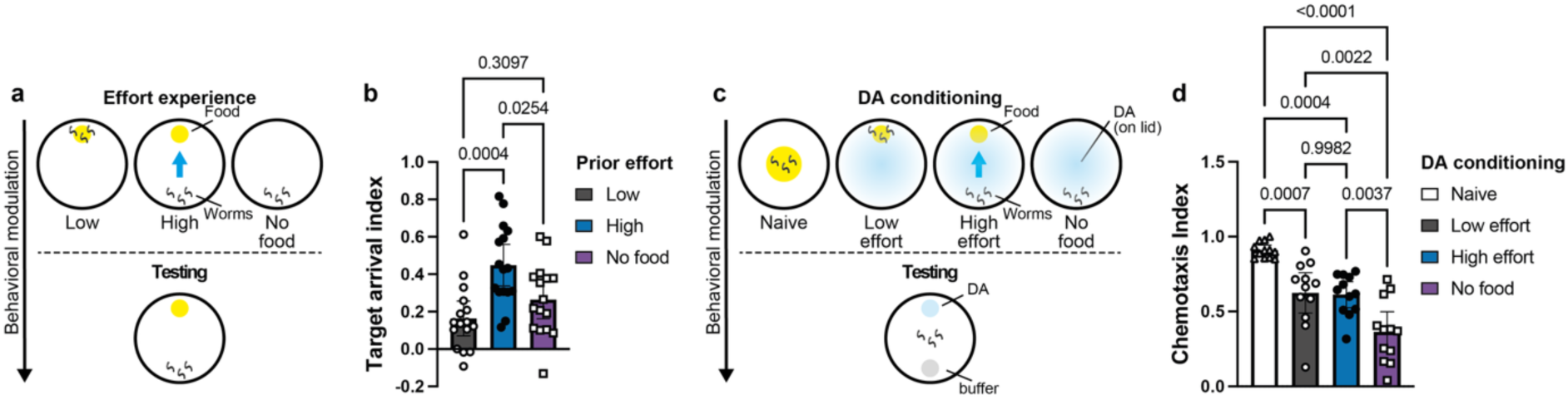
Food deprivation does not account for effort-induced behavioral modulation. (**a**) Schematic of low effort, high effort, and no-food experience followed by the food-approach test. (**b**) Target arrival following the indicated prior experiences. One-way ANOVA (*p* = 0.0005) followed by Tukey’s multiple comparisons test; *n* = 16 for each condition. (**c**) Schematic of diacetyl (DA) conditioning following the indicated prior experiences. During conditioning, a 5 μL drop of undiluted diacetyl was placed at the center of the plate lid. (**d**) Chemotaxis to DA following conditioning. One-way ANOVA (*p* < 0.0001) followed by Tukey’s multiple comparisons test; *n* = 12 for all conditions. Each point represents an independent experimental replicate (test plate). Bars show mean ± 95% CI.

Food deprivation can also act as an unconditioned stimulus in *C. elegans* associative learning, where pairing an attractive odor with the absence of food can reduce subsequent attraction to that odor. We therefore reasoned that, if the no food condition in our paradigm induced a meaningful deprivation state, it should be capable of driving this established form of behavioral plasticity. Animals underwent the same low effort, high effort, or no food experiences in the presence of the attractant diacetyl (DA), and their subsequent chemotaxis to DA was compared with that of naïve animals (Fig. 2c). Conditioning reduced attraction to DA under all three experience conditions relative to the naïve control, but the reduction was strongest following complete food deprivation, whereas low and high effort animals did not differ from one another (Fig. 2d). Thus, complete food deprivation was behaviorally effective, but the lack of any difference between low and high effort animals in this assay indicates that food deprivation is unlikely to account for the behavioral distinction between these conditions. The reduced attraction in the low and high effort conditions could be due to shared differences in handling or in the amount of food relative to the naïve condition, as well as non-associative olfactory adaptation^11^.

### Behavioral consequences of effort experience

Having established that prior effort alters subsequent food approach, we next altered the testing assays to ask how this modulation is expressed across different behavioral contexts. We first varied the demands of the food-approach test itself. In a less demanding version of the test, animals were placed at the middle of the plate rather than at the opposite edge, reducing the distance required to reach the food target (Fig. 3a). Conversely, we increased test demand by introducing an aversive barrier between the animals and the opposite side of the plate, either with food present beyond the barrier or with no food present (Fig. 3a). Across these conditions, prior high effort experience was associated with greater target arrival relative to low effort experience, despite considerable changes in performance of all worms (Fig. 3b). The persistence of the effect when animals were required to cross an aversive barrier, even in the absence of food, indicated that the post-effort modulation was not restricted to enhanced food attraction and was therefore unlikely to reflect a simple form of chemotaxis learning.

**Figure 3.**
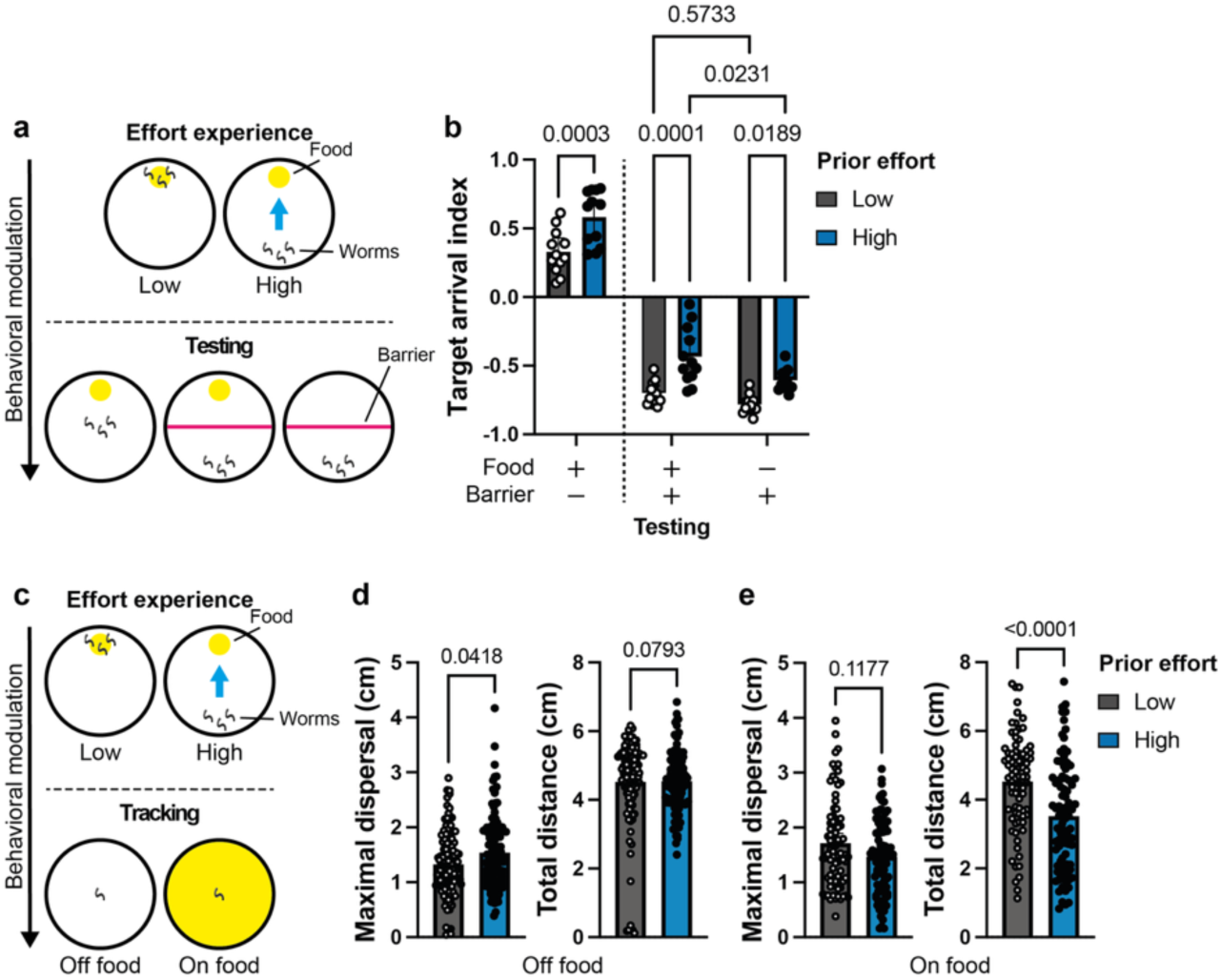
Behavioral consequences of effort experience. (**a**) Schematic of low and high effort experience followed by an easier food-approach test starting from the middle of the plate, or testing with an aversive barrier with or without food beyond it. (**b**) Target arrival under the indicated testing conditions. Two-way ANOVA (main effects of effort level and test demand, both *p* < 0.0001; interaction, *p* = 0.5139), followed by Šidak’s multiple comparisons test; *n* = 12 for each condition. (**c**) Schematic of low and high effort experience followed by short-term locomotion tracking off or on food. (**d**) Maximal dispersal and total distance traveled during 2 min of tracking off food. Mann–Whitney tests; *n* = 122, 117 (low, high). (**e**) Maximal dispersal and total distance traveled during 2 min of tracking on food. Mann–Whitney tests; *n* = 82, 90 (low, high). Each point represents an independent experimental replicate (test plate) in (b) and an individual tracked animal in (d,e). Bars show mean ± 95% CI.

We next asked whether this modulation could be detected in more basic features of locomotor behavior. Animals were tracked for 2 min either off food or on food (Fig. 3c). Off food, prior high effort increased maximal dispersal, whereas the total distance traveled was not significantly changed (Fig. 3d). Thus, high-effort animals did not simply move more, but their locomotion carried them farther from their starting position. On food, by contrast, prior high effort markedly reduced the total distance traveled, while maximal dispersal showed no significant change (Fig. 3e). Together, these results indicate that effort experience alters subsequent locomotion in a context-dependent manner: in the absence of food, movement becomes more spatially dispersive without an overall increase in locomotor output, whereas after food encounter locomotor activity is suppressed.

### Temporal persistence of effort-induced behavioral modulation

Having found that effort experience leaves a measurable trace in subsequent behavior, we next asked whether this trace persists beyond the immediate post-effort period, which would distinguish it from a short-lived carryover of arousal or locomotor activation. To address this, animals were transferred after the effort experience to a fresh plate containing food, and allowed to remain there before testing (Fig. 4a). The difference between low and high effort animals was still evident after 1 h on food, with high effort animals retaining enhanced target arrival relative to low effort controls (Fig. 4b). Thus, the behavioral effect outlasted the effort episode itself and persisted despite an extended period of feeding and recovery, arguing against a short-lived spillover of behavioral activation. Persistence during this period on fresh food also further weakens food deprivation as an explanation for the altered behavior.

**Figure 4.**
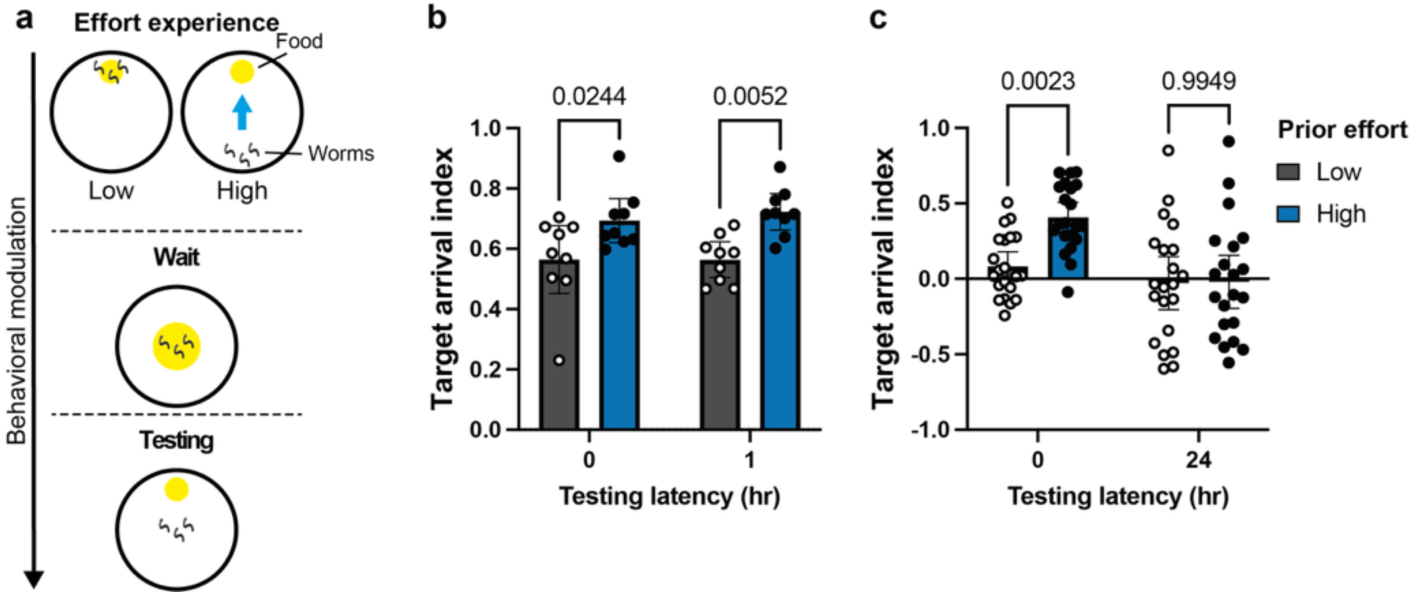
Effort-induced behavioral modulation persists for 1 h but is lost after 24. **h**. (**a**) Schematic of low and high effort experience followed by a waiting period on food before subsequent food-approach testing. (**b**) Target arrival index immediately or 1 h after effort experience, using the mid-plate food-approach test. Two-way ANOVA (main effect of effort level, *p* = 0.0002; main effect of latency, *p* = 0.6774; interaction, *p* = 0.6695), followed by Šídák’s multiple comparisons test; *n* = 9 for each condition. (**c**) Target arrival index immediately or 24 h after effort experience, using the full-plate food-approach test. Two-way ANOVA (main effect of effort level, *p* = 0.0167; main effect of latency, *p* = 0.0002; interaction, *p* = 0.0230), followed by Šídák’s multiple comparisons test; *n* = 21 for each condition. Each point represents an independent experimental replicate (test plate). Bars show mean ± 95% CI.

We next extended the interval between effort experience and testing to 24 h. In contrast to the 1 h condition, the difference between low and high effort animals was no longer detectable after 24 h (Fig. 4c). The modulation therefore has a finite lifetime, present at 1 h and absent by 24 h. Loss of the effect after 24 h also argues against the possibility that the high effort procedure simply selected a stable subpopulation of particularly fast or effective animals when worms were recovered from the food patch. Such selection would be expected to produce a persistent difference between the groups rather than one that disappears with time. Moreover, most high effort animals reached the food patch during the 1.5 h experience period, further limiting the scope for such selection. Together, these results indicate that effort experience produces a persistent but time-limited behavioral trace.

### Tyramine signaling contributes to effort-induced behavioral modulation

Monoamines are major regulators of behavioral state, motivation, arousal, and locomotion across animals, and in *C. elegans* they strongly influence foraging, food-dependent behavior, and experience-dependent plasticity. Because effort experience combines sustained locomotion, food seeking, and a subsequent change in behavioral state, we asked whether monoaminergic signaling contributes to the resulting modulation. We first tested *cat-1* mutants, which lack the vesicular monoamine transporter required for normal monoamine storage and release^12^. Loss of *cat-1* substantially altered performance after both low and high effort experience and, importantly, reduced behavioral modulation, as measured by the difference between the target arrival indices of these two conditions (Fig. 5b). This pointed to a role for monoaminergic signaling in the generation, maintenance, or expression of effort-driven behavioral modulation.

**Figure 5.**
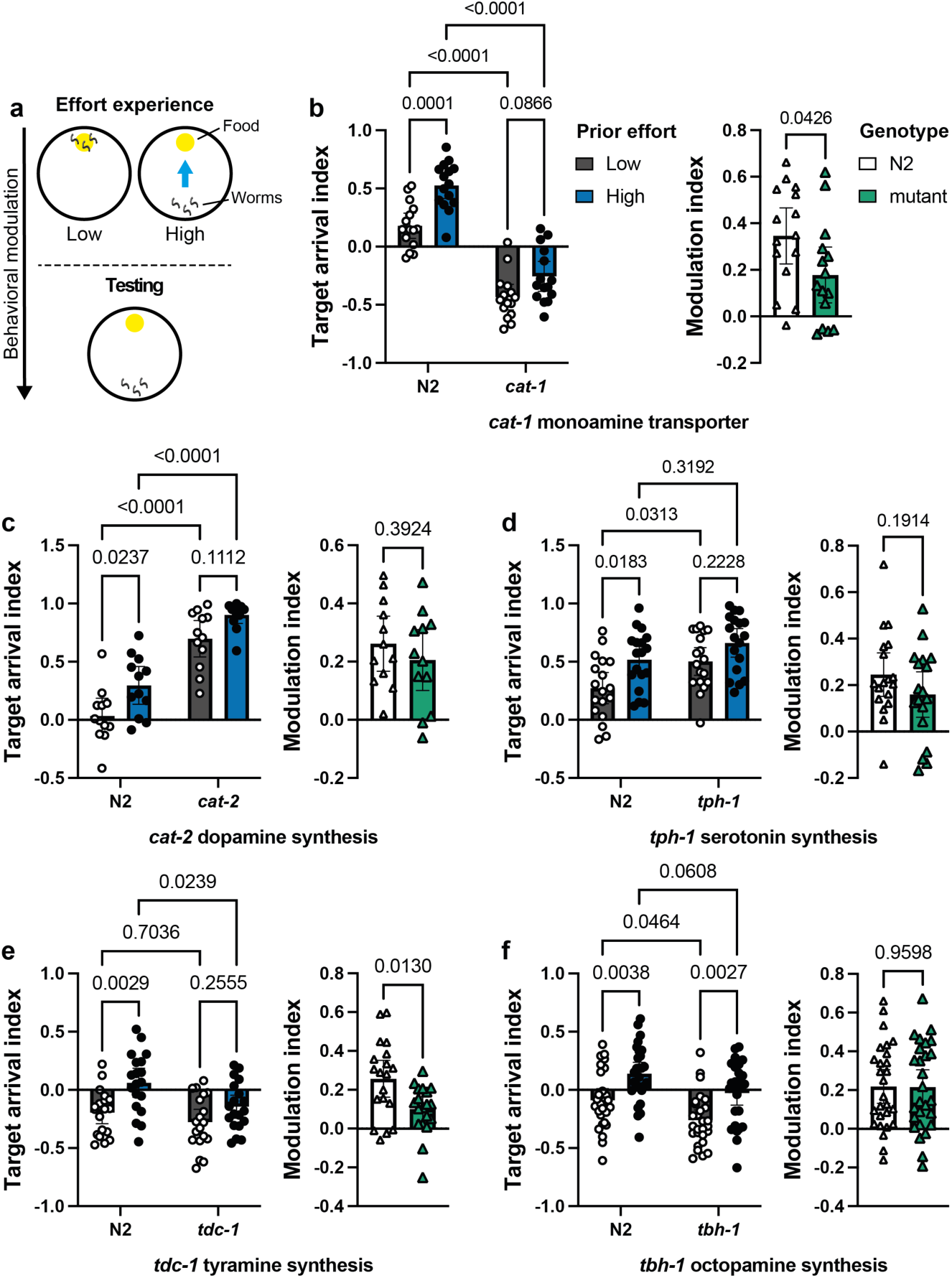
Tyramine signaling contributes to effort-induced behavioral modulation. (**a**) Schematic of low and high effort experience followed by food-approach testing. (**b–f**) Target arrival indices of N2 and the indicated monoamine-related mutants following low or high effort experience (left), and corresponding behavioral modulation (high − daily low average; right). Arrival indices were analyzed using two-way ANOVA followed by Šídák’s multiple-comparisons test. ANOVA *p*-values for genotype, effort, and their interaction, respectively: (**b**) *cat-1*, <0.0001, <0.0001, 0.1219; (**c**) *cat-2*, <0.0001, 0.0007, 0.6574; (**d**) *tph-1*, 0.0025, 0.0011, 0.4711; (**e**) *tdc-1*, 0.0066, 0.0003, 0.2333; (**f**) *tbh-1*, 0.0006, <0.0001, 0.9416. Modulation indices were analyzed using unpaired t-tests with Welch’s correction (b–d, f) or a Mann–Whitney U test (e). (**b**) *n* = 15; (**c**) *n* = 12; (**d**) *n* = 18; (**e**) *n* = 19; (**f**) *n* = 27. Each point represents an independent experimental replicate. Bars show mean ± 95% CI.

We therefore examined individual monoamine biosynthetic pathways. We first examined *cat-2*, which encodes tyrosine hydroxylase, the rate-limiting enzyme in dopamine synthesis^13^. *cat-2* mutants showed strongly elevated performance relative to N2 following both low and high effort experience, while retaining a similar level of effort-dependent behavioral modulation as N2 (Fig. 5c). Dopamine signaling therefore had a substantial influence on overall performance in the assay but was not specifically required for effort-induced modulation. We next examined *tph-1*, which encodes tryptophan hydroxylase, the rate-limiting enzyme in neuronal serotonin synthesis^14^. Loss of *tph-1* likewise shifted performance under both experience conditions, while producing only a comparatively small reduction in effort-dependent modulation (Fig. 5d). Thus, although both dopamine and serotonin substantially shaped behavior in the assay, neither mutant reproduced the selective disruption of effort-dependent modulation suggested by *cat-1*.

A more specific effect emerged from the tyramine pathway. *tdc-1* encodes tyrosine decarboxylase, which converts tyrosine to tyramine and thereby also supplies the precursor for octopamine synthesis^15^. In *tdc-1* mutants, performance following high effort was selectively reduced relative to N2, resulting in a significant attenuation of effort-induced modulation (Fig. 5e). This phenotype could reflect loss of tyramine itself or loss of octopamine synthesized downstream from tyramine. We therefore examined *tbh-1* mutants, which lack tyramine β-hydroxylase and are unable to convert tyramine to octopamine^15^. In contrast to *tdc-1*, *tbh-1* animals retained effort-induced modulation (Fig. 5f), pointing to tyramine rather than octopamine as the more likely contributor.

These results identify tyramine signaling as a candidate component of the system linking effort experience to subsequent behavioral change. Tyramine could participate in generating or encoding the effort experience itself, in maintaining the resulting behavioral state, in modulating behavior during the subsequent test, or at more than one of these stages. Distinguishing among these possibilities will require temporally and spatially restricted manipulation of tyraminergic signaling.

### Effort experience modulates RIM and AWC neuronal activity

The genetic results pointed to tyramine as a candidate mediator of effort-induced behavioral modulation, directing our attention to the RIM interneurons, a principal tyraminergic source in the *C. elegans* nervous system^15^. RIM plays an important role in locomotor-state control, particularly in reversals and transitions between movement states^16^, and participates in circuit plasticity underlying learned behavioral changes^17^. We therefore asked whether effort experience leaves a detectable trace in ongoing RIM activity. Calcium imaging following low or high effort experience revealed reduced spontaneous RIM activity after high effort (Fig. 6a,b). Both the number of calcium transients and their average amplitude were reduced (Fig. 6c), indicating that effort experience is accompanied by a subsequent change in RIM activity.

**Figure 6.**
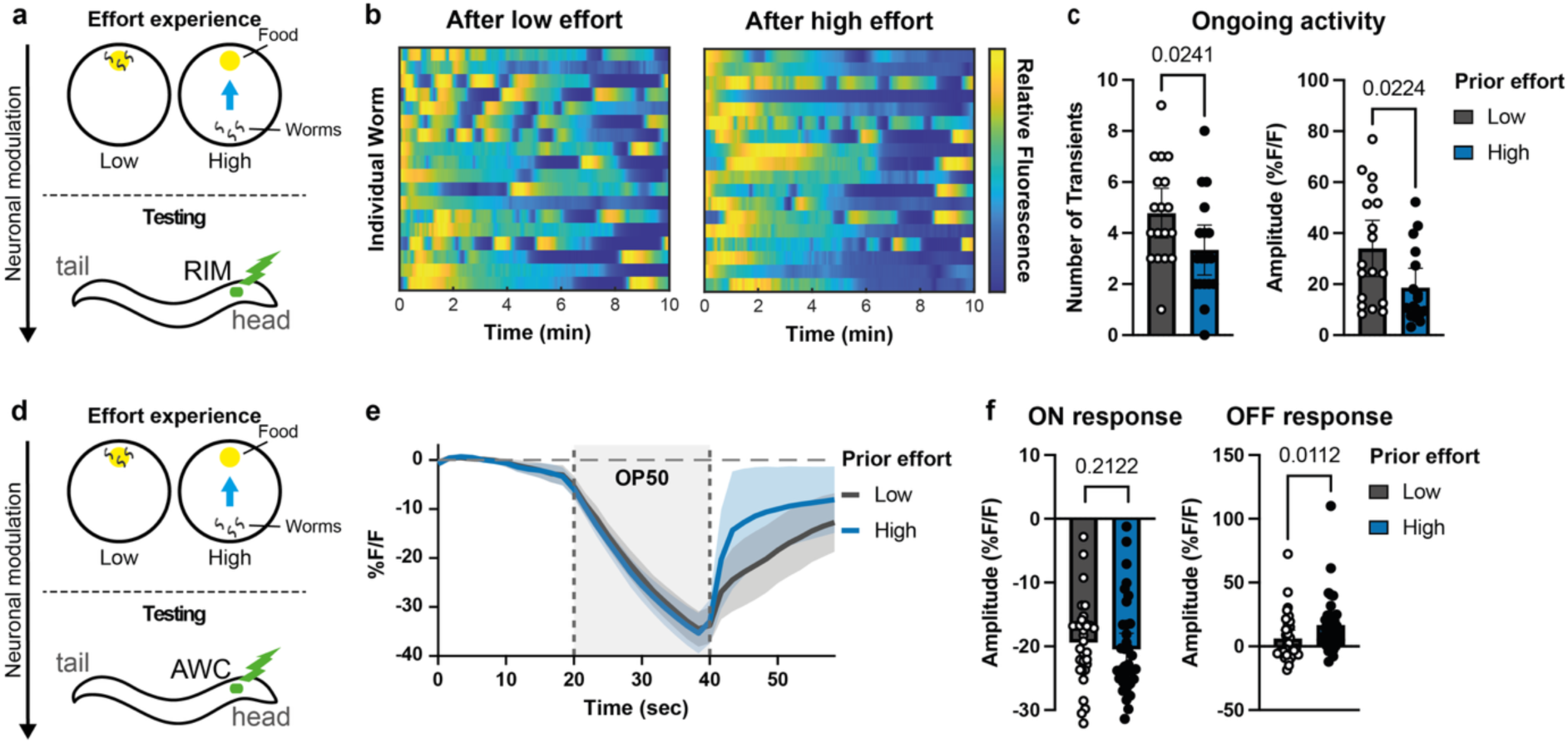
Effort experience alters RIM and AWC neuronal activity. (**a**) Schematic of low and high effort experience followed by calcium imaging of RIM. (**b**) Heat maps showing normalized RIM fluorescence over 10 min in individual animals following low or high effort experience. (**c**) Quantification of ongoing RIM activity, measured as the number of calcium transients and their mean amplitude. Mann-Whitney tests; left, *n* = 18 for each condition; right, *n* = 18, 17 (low, high). (**d**) Schematic of low and high effort experience followed by calcium imaging of AWC. (**e**) Mean AWC fluorescence traces during odor presentation and removal following low or high effort experience. Shaded bands show 95% CI. Vertical dashed lines indicate OP50 bacteria presentation and removal. (**f**) Quantification of AWC ON and OFF response amplitudes. Mann-Whitney tests; *n* = 33, 35 (low, high) in both left and right. Each point represents an individual animal. Bars show mean ± 95% CI.

This result was reminiscent of our previous findings in *mec-4* mutants, in which ongoing RIM activity is also reduced^18^. The associated locomotor phenotypes, however, differ in important ways. *mec-4* animals show reduced reversal rates on food^19^, but increased reversal rates and reduced dispersal off food^18^. Following high effort, by contrast, off-food locomotion becomes more spatially dispersive without a corresponding increase in total distance traveled, whereas on food overall locomotor output is reduced. Thus, a similar reduction in ongoing RIM activity can accompany distinct, and in some respects opposing, reorganizations of context-dependent locomotor behavior.

We therefore asked whether AWC sensory responses might distinguish these states. In *mec-4* mutants, AWC responses themselves were unchanged, even at very low odor concentrations^19^, whereas the downstream AIY response to removal of an AWC-sensed odor was altered, showing an enhanced inhibitory OFF response^19^. Following effort experience, by contrast, AWC responses were themselves modified: the response to odor onset was unchanged, whereas the OFF response following odor removal was enhanced after high effort (Fig. 6d–f). This suggests that the post-effort state differs from the *mec-4* state not only in its behavioral expression, but also in the locus of sensory-circuit modulation.

Together, these findings identify RIM and AWC as neuronal correlates of the post-effort state. Reduced RIM activity may represent one component shared with other locomotor states, while the accompanying change in AWC sensory responses may help distinguish the specific behavioral consequences of effort experience. At present, however, the causal relationships among tyramine signaling, RIM activity, AWC modulation, and the behavioral phenotype remain to be resolved.

## Discussion

Here we show that a relatively simple increase in the effort required to obtain food can leave a persistent trace that alters subsequent behavior in *C. elegans*. The effect was distinct from the consequences of food deprivation, was generalized across several test formats, and persisted for at least 1 h. Its behavioral expression extended beyond enhanced food approach, including increased barrier crossing even in the absence of food, while locomotor tracking revealed context-dependent changes in spatial dispersal and locomotor output. These observations suggest that effort itself can act as a salient component of experience, capable of shaping subsequent behavioral state.

### Ordinary effort as a source of persistent behavioral modulation

Effort accompanies many ordinary behaviors, but it is often studied in situations where an animal chooses between actions that differ in their costs and expected outcomes, linking effort closely to valuation, motivation, and action selection^6,7^, recently also in *C. elegans*^10^. Our paradigm addresses a complementary form of effort: one imposed by the structure of the environment^4,5^, in which the eventual outcome is held constant while the behavioral demand required to attain it varies. Although the paradigm may superficially resemble operant conditioning, both low and high effort animals ultimately receive the same food outcome, and the critical manipulation is the amount of work required to reach it rather than whether a particular action is selectively reinforced. We therefore use effort operationally to describe the greater behavioral demand associated with attaining the same outcome, without implying subjective choice or formally defined goal-directed action.

Previous work has shown that imposed effort can influence subsequent persistence and performance^4,5^. Our findings extend this idea in several ways. Here, the effort is embedded in an ordinary and recurrent behavior, approaching food, rather than in an explicitly demanding task. Its consequences are also broader than a simple increase in persistence or behavioral vigor: prior effort altered performance across distinct test formats and changed how locomotion was organized across food contexts. Off food, high effort animals achieved greater maximal dispersal without increasing total distance traveled, whereas on food they showed a marked reduction in total distance traveled. The effect furthermore persisted beyond the original episode, indicating that even modest variation in the demands encountered during routine behavior can leave a temporally extended behavioral trace.

### A minimal form of experience-dependent behavioral plasticity

Many forms of learning involve modifying the significance of a particular stimulus, action, or outcome. Classical conditioning links a cue to an appetitive or aversive event, while other paradigms alter behavior through reinforcement, repeated exposure, or changes in stimulus contingencies. Here, the relevant experience is embedded in a basic behavior, approaching food, and does not require assigning new value to a discrete object or sensory cue. The subsequent behavioral change is likewise not restricted to altered responses toward a particular object: although food approach is modified, the effect also generalizes to other test contexts, arguing against a simple form of chemotaxis learning.

These observations raise the possibility that experience-dependent modulation is continuously incorporated within normal behavior, recursively shaping future behavior. Such recursion could in principle lead to cumulative behavioral divergence, as each action alters the state from which the next action emerges. Alternatively, behavioral systems may be organized around a more stable baseline, with recent experience producing temporary deviations that are subsequently erased through forgetting or other restorative processes. The disappearance of the effort-induced effect by 24 h is consistent with such recentering, although the mechanisms underlying this return remain unknown.

### Tyramine and circuit-state modulation after effort

Tyramine emerged as a candidate signal linking effort experience to subsequent behavioral change. Its involvement raises a fundamental question about how the effort episode is represented over time: tyramine could contribute to encoding the experience as it occurs, to maintaining the resulting state, or to expressing that state during subsequent behavior. These possibilities place tyraminergic signaling at a potential interface between the experience of effort and its later behavioral consequences.

RIM, the principal tyraminergic neuron class in *C. elegans*^15^, provides a plausible neuronal counterpart to this signal. It is particularly well positioned to couple neuromodulatory state to locomotor circuitry: tyramine signaling from RIM is predominantly extrasynaptic and reaches a broad receptor-defined network, and RIM ranks among the neurons with the strongest participation across the monoaminergic, synaptic, and gap-junction connectomes^20^. We found that high effort reduced ongoing RIM activity, consistent with a shift in the internal locomotor state of the animal. As noted above, a similar reduction in RIM activity has been associated with cross-modal plasticity in *mec-4* mutants^18^, which lack gentle-touch sensation^21^. Yet the resulting locomotor reorganization differs markedly from that observed after effort. Thus, reduced ongoing RIM activity appears compatible with distinct context-dependent behavioral states rather than specifying a single locomotor outcome.

The AWC results suggest one possible basis for this divergence. In *mec-4* mutants, AWC sensory responses themselves remain largely unchanged, while altered processing appears downstream at AIY^19^. Following high effort, by contrast, the AWC OFF response itself is enhanced. A shared reduction in RIM activity may therefore be embedded within different sensory-circuit configurations, helping to explain why the two forms of plasticity produce distinct food-dependent locomotor states.

Tyramine also provides an interesting comparative link beyond *C. elegans*. Across invertebrates, tyramine functions as an independent neuromodulator, contributing to the regulation of behavior, metabolism, and physiological state^22,23^. In insects, tyramine can act independently of, and sometimes antagonistically to, octopamine. For example, the two amines exert opposing effects on locomotion and its adaptation to nutritional state in *Drosophila*^24,25^. Octopamine itself is often considered a functional analogue of vertebrate noradrenaline^22^, placing tyramine within an aminergic system with broad roles in behavioral-state regulation. In vertebrates, tyramine persists at low concentrations as a trace amine, where its physiological roles remain incompletely resolved but include signaling through TAAR1 and modulation of classical monoaminergic systems^26^. Thus, although the evolutionary and functional correspondences are not one-to-one, the involvement of tyramine raises the possibility that effort-dependent behavioral modulation draws on an evolutionarily conserved aminergic logic for regulating behavioral state.

Together, our findings position effort as a feature of experience that can be retained beyond the action in which it arises, integrated with current sensory context, and translated into subsequent behavioral change. The involvement of tyramine, together with post-effort changes in RIM and AWC activity, provides an initial circuit-level framework for understanding how such experience is encoded and later expressed.

More broadly, this framework offers a tractable route toward elucidating how the nervous system continuously incorporates an animal’s own recent behavior into the state from which future behavior emerges.

## Materials and Methods

### *C. elegans* growth and maintenance

*C. elegans* strains were maintained under standard conditions at room temperature (20-21°C) on nematode growth medium (NGM) 2% agar plates seeded with *E. coli* strain, OP50. The N2 strain (Bristol, England) was used as the reference strain. Other strains used in the study are listed in Table 1.

**Table 1.**
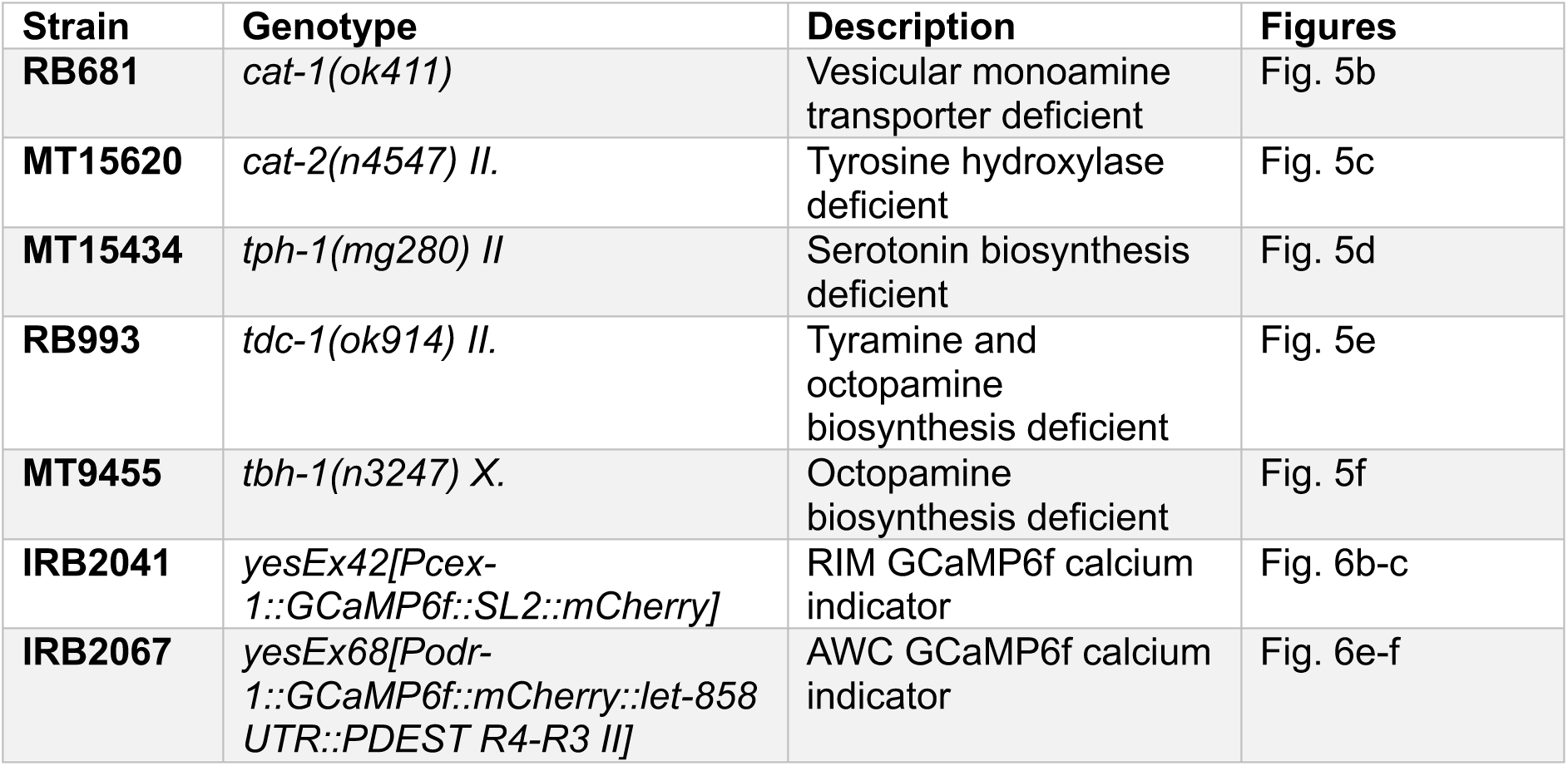
List of C. elegans strains used in this study.

| Strain | Genotype | Description | Figures |
| --- | --- | --- | --- |
| **RB681** | *cat-1(ok411)* | Vesicular monoamine transporter deficient | Fig. 5b |
| **MT15620** | *cat-2(n4547) II.* | Tyrosine hydroxylase deficient | Fig. 5c |
| **MT15434** | *tph-1(mg280) II* | Serotonin biosynthesis deficient | Fig. 5d |
| **RB993** | *tdc-1(ok914) II.* | Tyramine and octopamine biosynthesis deficient | Fig. 5e |
| **MT9455** | *tbh-1(n3247) X.* | Octopamine biosynthesis deficient | Fig. 5f |
| **IRB2041** | *yesEx42[Pcex-1::GCaMP6f::SL2::mCherry]* | RIM GCaMP6f calcium indicator | Fig. 6b-c |
| **IRB2067** | *yesEx68[Podr-1::GCaMP6f::mCherry::let-858 UTR::PDEST R4-R3 II]* | AWC GCaMP6f calcium indicator | Fig. 6e-f |

### Effort training paradigm

Effort training was performed on 90 mm NGM plates. One day prior to the assay, two marks were made on opposite sides of each plate, 13 mm from the edge. A 100 µL drop of OP50 (OD600 = 1.0) was placed on one of the marks and allowed to dry for 18– 22 h. On the day of the assay, synchronized young adult worms were washed from their growth plates using M9 buffer and subsequently washed three additional times with M9 to remove residual bacteria. Washed worms were then transferred either directly onto the food source (low effort condition) or onto the mark on the opposite side of the plate (high effort condition). Excess M9 was removed using Kimtech wipes, and worms were allowed to remain under the respective conditions for 1.5 h. Following training, worms located on the food source were collected and washed prior to preparation for subsequent assays. For locomotion and calcium imaging experiments, in which individual animals were analyzed, worms were individually picked rather than collected by washing.

For experiments including a no-food condition, worms were subjected to the same handling procedure but were placed on one mark of an empty NGM plate that had been maintained at room temperature on the laboratory bench for one day prior to the assay, consistent with the preparation of plates used for the high and low effort conditions. Because worms in the no-food condition were collected from the entire plate, worms from all other experimental conditions in the same experiment were likewise collected from the entire plate rather than exclusively from the food source.

### Food arrival assay

Following effort training, arrival at OP50 was assessed on 90 mm NGM plates. One day prior to the assay, two marks were made on opposite sides of each test plate, 13 mm from the edge, and a line was drawn across the middle of the plate. A 30 µL drop of OP50 (OD600 = 1.0) was placed on one of the marks and allowed to dry for 18–22 h. Ten minutes prior to the end of the training period, 1 µL of 1 M sodium azide was added to the OP50 spot. At the end of the training period, worms were collected and washed three times with M9 buffer to remove residual bacteria. Washed worms were transferred to the mark opposite the OP50 spot, excess M9 was removed using Kimtech wipes, and worms were allowed to chemotax for 20 min. Following the assay, plates were placed at −20°C for 12 min to immobilize the worms and subsequently transferred to 4°C until scoring.

Worms that had crossed the middle line into the target half of the plate were counted and compared with worms remaining in the non-target half. Worms visibly remaining at the starting point, typically appearing as a small cluster of worms that has not dispersed from the transfer location, were excluded from the analysis as they may have been injured during transfer and consequently failed to move. The target arrival index was calculated as (number of worms in the target half − number of worms in the non-target half) / total number of worms. For mutant analyses, N2 and mutant conditions were balanced within experimental day and each high effort replicate was referenced to the corresponding same-day low effort mean when deriving the modulation index.

The easier version of the assay was similar to the standard, except that a third mark was made at the center of the plate as well as two lines 10 mm apart marking the boundaries of the starting zone. Animals were placed at the center of the plate rather than at the edge opposite the food source. OP50 was placed on one side of the plate and double-distilled water (DDW) on the opposite side. Ten minutes prior to the end of the training period, 1 µL of 1 M sodium azide was added to both the OP50 and DDW spots. Following the 20-min chemotaxis assay, worms on the OP50 and DDW sides were counted separately. Worms remaining between the two central lines, including those at the starting point, were excluded from the analysis. The chemotaxis index was calculated per plate as (number of worms on the OP50 side − number of worms on the DDW side) / total number of worms counted on either side.

### Aversive conditioning assay

Negative associative learning was assessed by conditioning worms with diacetyl during the training period, followed by measurement of their subsequent chemotaxis toward diacetyl using a protocol adapted from^27,28^. During conditioning, low effort, high effort, and no food groups were exposed to 5 µL of 100% diacetyl placed in the center of the plate lid. Plates were wrapped with Parafilm to allow a diacetyl gradient to form and persist throughout the 1.5-h conditioning period. Following conditioning, the different experimental groups, including a naïve group consisting of worms washed directly from their growth plates, were tested for chemotaxis toward diacetyl.

Chemotaxis assay plates were prepared 18–22 h prior to the assay and maintained at room temperature on the laboratory bench. Three marks were made on each plate: two marks 13 mm from opposite edges of the plate,one mark at the center and two lines 10 mm apart marking the boundaries of the starting zone. Ten minutes prior to the end of the conditioning period, 1 µL of 1 M sodium azide was added to each of the two side marks. At the end of the conditioning period, worms were collected and washed three times with M9 buffer to remove residual bacteria and transferred to the center mark. A 3-µL drop of 1:10,000 diacetyl, diluted in 100% ethanol, was placed on one side mark, while 3 µL of 100% ethanol was placed on the opposite side mark. Worms were allowed to crawl for 45 min. Following the assay, plates were transferred to −20°C to immobilize the worms and subsequently transferred to 4°C until scoring. Worms on the diacetyl and ethanol sides were counted separately. Worms remaining between the two lines around the center mark, including those at the starting point, were excluded from the analysis. The chemotaxis index was calculated per plate as (number of worms on the diacetyl side − number of worms on the ethanol side) / total number of worms counted on either side.

### Glycerol barrier-crossing

Following effort training, glycerol barrier-crossing behavior was assessed on 90-mm NGM plates using an assay adapted from previously described methods^29^. One day prior to the assay, two marks were made on opposite sides of each plate, 13 mm from the edge, and a line was drawn across the middle of the plate. A 30-µL drop of OP50 (OD600 = 1.0) was placed on one of the marks and allowed to dry for 18–22 h. In experiments with no food target this step was omitted. Plates were maintained at room temperature on the bench. In experiments including the food target, 1 µL of 1 M sodium azide was added to the OP50 spot 10 min prior to the end of the training period. Five minutes prior to the end of the training period, 30 µL of 4.5 M glycerol was applied along the middle line and spread across the line using a thin brush.

At the end of the training period, worms were collected and washed three times with M9 buffer to remove residual bacteria. Worms were then transferred to the mark opposite the OP50 patch, when present, or to a side mark in the absence of a food target. Excess M9 was removed using Kimtech wipes, and worms were allowed to roam freely for 20 min. Following the assay, plates were placed at −20°C for 12 min to immobilize the worms and subsequently transferred to 4°C until scoring.

Worms that had crossed the glycerol barrier line into the target half of the plate were counted and compared with worms remaining in the non-target half. Worms remaining at the starting point were excluded from the analysis, as they may have been injured during transfer and consequently failed to move. The target arrival index was calculated as (number of worms in the target half − number of worms in the non-target half) / total number of worms included in the analysis.

### Memory assay

To assess memory retention following effort training, worms were washed with M9 and transferred to fresh OP50-seeded NGM plates for a defined waiting period before the food chemotaxis assay. For the 1 h retention condition, worms were transferred to plates seeded with 500 µL OP50 grown overnight at 37°C. For the 24 h retention condition, worms were transferred to plates seeded with 1 mL OP50 grown overnight at 37°C. Following the respective waiting period, worms were tested for chemotaxis toward OP50 as described above.

### Locomotion tracking and analysis

Following effort training, 5–10 worms were transferred to 50 mm test plates for locomotion tracking. For tracking on food, plates were seeded with 200 µL OP50 and incubated at 37°C for 1–2 h, followed by 1 h at room temperature. Worms were transferred first to an empty plate for 1 min to remove residual bacteria and then to the test plate, where they were allowed to acclimate for 1 min before 2-min recordings at 26.1 frames/s using a Pixelink camera. For tracking off food, unseeded plates were dried with the lid open in a laminar-flow hood for 1 h before the assay, and worms were recorded for 2 min following the same transfer and acclimation procedure.

Videos were analyzed using the TReX animal tracking system^30^, and X–Y centroid coordinates were extracted for each worm. Trajectories were analyzed in MATLAB to calculate total distance and maximum dispersal. Total distance was calculated as the cumulative Euclidean distance between consecutive positions and maximum dispersal as the maximum Euclidean distance from the starting position.

### Calcium Imaging of Neuronal Activity

Following effort training, calcium imaging was performed to assess spontaneous activity in RIM and stimulus-evoked activity in AWC. Imaging was performed using microfluidic devices designed for each experimental paradigm. RIM spontaneous activity was recorded using a PDMS microfluidic chip that confines the worm’s head while allowing body movement, as previously described^31^. The chip was mounted on a Zeiss AxioObserver 7 inverted fluorescence microscope with a 40× air objective and a Photometrics Prime 95B sCMOS camera. Worms were immobilized in M9 containing 10 mM levamisole and recorded for 10 min. AWC stimulus-evoked activity was recorded using an olfactory evoked-activity chip that enabled controlled delivery of OP50 (OD600 = 1.0) to the worm’s nose^31^. Worms were immobilized in 5 mM levamisole diluted in DDW. OP50 was supplemented with rhodamine (0.0002 mM) to verify stimulus delivery. The stimulus protocol consisted of 1 min of blue-light illumination for adaptation, followed by 20 s DDW, 20 s OP50, and 20 s DDW.

For both experiments, mean raw fluorescence within the neuronal ROI was extracted using a custom-written script and analyzed using a MATLAB program as previously described^19^. For RIM, the amplitude and number of spontaneous calcium transients were quantified. For AWC, stimulus responses were quantified by averaging fluorescence over 10 frames immediately before and after stimulus switch. The stimulus difference was calculated as the percentage change in fluorescence relative to the pre-stimulus baseline, (F_after – F_before)/ F_before*100.

### Statistical analysis

Statistical analysis was performed using Prism GraphPad (Version 11.1.0). For each dataset, initial normality tests were carried out in order to determine whether to perform a parametric or non-parametric test. Details about sample sizes and the statistical tests applied are provided in the figure legends. Raw data are provided in the Supporting Data File.

## Acknowledgements

This work was supported by Israel Science Foundation Grant 1465/20 and by the Horizon Europe, PathFinder European Innovation Council Work Programme under grant agreement No 101098722. Views and opinions expressed in this article are those of the authors only and do not necessarily reflect those of the European Union or European Innovation Council and SMEs Executive Agency (EISMEA). Neither the European Union nor the granting authority can be held responsible for them. We are grateful to the *Caenorhabditis elegans* Genetic Consortium (funded by NIH Office of Research Infrastructure Programs P40 OD010440).

